# Glutamate and early functional NMDA Receptors promote axonal elongation modulating both actin cytoskeleton dynamics and H_2_O_2_ production dependent on Rac1 activity

**DOI:** 10.1101/2025.02.18.638875

**Authors:** Ernesto Muñoz-Palma, Carlos Wilson, Cecilia Hidalgo, Christian González-Billault

## Abstract

NMDA Receptors (NMDARs) have essential functions in the nervous system, including neuronal maturation, neurotransmission, synaptic plasticity, learning, and memory. Following membrane depolarization and glutamate activation, NMDARs mediate Ca^2+^ influx into neurons, activating Ca^2+^ signaling cascades with key roles in neuronal function. However, no studies have been reported on the roles of glutamate and NMDARs during early neuronal development. Although NMDARs classically act at the postsynaptic membrane, the present results indicate that neurons express functional NMDARs during polarity acquisition and localize them in the axonal compartment early in development; at this stage, cultured neurons spontaneously release glutamate. In addition, pharmacological and genetic experiments for NMDARs loss- and gain-of-function modulated neuronal polarization and axonal elongation antagonistically. An intracellular mechanism involving Ca^2+^ release from the endoplasmic reticulum, activation of the Rho GTPase Rac1, and actin cytoskeleton rearrangements at the axonal growth cone couples these morphological changes. Moreover, NMDAR activity regulates the physiological intracellular production of hydrogen peroxide (H_2_O_2_) via a Rac1/NADPH oxidase complex to support neuronal development. Optogenetic Rac1 activation simultaneously promoted lamellipodia formation and H_2_O_2_ production suggesting functional coupling between these seemingly unconnected events. The mechanism presented here involves a dual function for the Rac1 protein that depends on glutamate and NMDAR activity. Based on these findings, we suggest that early physiological and spontaneous glutamate release activates NMDARs to promote early neuronal development before synapse formation, indicating that glutamate is necessary for neurotransmission, early neuronal development, and axonal growth.

## Introduction

The N-methyl-D-aspartate receptors (NMDARs) are Ca^2+^ channels activated by glutamate, the principal excitatory neurotransmitter in the nervous system. NMDARs play roles in synaptic transmission, plasticity, learning, memory, and neurodevelopment (Traynelis et al., 2010; Paoletti, Bellone, and Zhou, 2013; Iacobucci and Popescu, 2017). These channels form a heterotetrameric structure typically comprised of two GluN1 and two GluN2 subunits (also known as NR1 and NR2, respectively) that allow Ca^2+^ influx when activated (Traynelis et al., 2010; Paoletti, Bellone, and Zhou, 2013). For proper connectivity and neurotransmission, neurons possess two structurally and functionally different compartments that emerge from the cell body (soma): a single axon and several dendrites to transmit and receive inputs, respectively. The highly polarized morphology of neurons and the axon specification and outgrowth are the result of a sequence of stages of molecular, cellular, and morphological changes that collectively are termed the establishment of neuronal polarity (Banker and Cowan, 1977; Dotti, Sullivan, and Banker, 1988; Cáceres, Ye and Dotti, 2012), which occurs during early neuronal development and before the formation of synaptic contacts. Thus, acquiring neuronal morphology is essential for proper connectivity and excitatory neurotransmission, mediated mainly by glutamate receptors in hippocampal neurons. The NMDARs are commonly distributed into dendrites, controlling their growth and dendritic spine maturation (Sepulveda et al., 2010; Bustos et al., 2014), and thus have a pivotal role in synaptic transmission (Daw, Stein and Fox, 1993).

However, a growing body of evidence shows that NMDARs also have a role in neuronal development. Previous reports indicate that NMDAR activity promotes the migration of embryonic cortical and cerebellar granule neurons during early mouse development (Komuro and Rakic, 1993; Behar et al., 1999; Kumada and Komuro, 2004). Experiments in cultured cerebellar granule neurons showed that NMDAR activation promotes neuritogenesis (Pearce, Cambray-Deakin, and Burgoyne, 1987; Rashid and Cambray-Deakin, 1992).

Moreover, previous studies in cultured Xenopus spinal neurons suggest that NMDARs mediate axonal growth cone turning (Zheng, Wan, and Poo, 1996). In cultured dopaminergic neurons, it has been suggested that NMDARs might be expressed at the axonal growth cones and that their activation could promote axonal elongation and growth cone splitting (Schmitz et al., 2009). More recently, it was reported that NMDARs are expressed in the axons of developing neocortical neurons (Gill et al., 2014) and are functional in the growth cone of already polarized neurons (Wang et al., 2011). However, the expression, localization, and role of NMDARS during the early stages of neuronal polarization remain unreported.

It has been widely described that the Rho GTPases family and the actin cytoskeleton are essential regulatory molecules involved in neuronal development (Gonzalez-Billault et al., 2012; Stankiewicz and Linseman, 2014). In addition, emerging reports suggest that Reactive Oxygen Species (ROS) production mediated by the NADPH oxidase 2 (NOX2 complex) promotes neuronal polarity and axonal outgrowth by a mechanism that involves Ca^2+^ signaling, Rho GTPases activity and actin cytoskeleton dynamics (Munnamalai and Suter, 2009; Munnamalai et al., 2014; Wilson et al., 2015; Wilson et al., 2016; Wilson et al., 2018). In addition, previous reports showed that abnormally high NMDAR stimulation promotes NOX2-dependent ROS production and cellular death in mature cortical neurons (Brennan et al., 2009; Reyes et al., 2012). Currently, the paradigm that ROS are only detrimental species is outdated, as the physiological production of ROS, termed oxidative eustress, is essential for the maintenance and proliferation of neural stem cells and for defining exquisite and complex neuronal morphology. Yet, elevated and accumulative ROS production, termed oxidative distress, is deleterious for neurons and promotes neuronal dysfunction and neurodegeneration (Sies and Jones, 2020). These divergent roles displayed by ROS emphasize the fact that more studies are needed to understand how they exert their different physiological and pathological effects on neuronal cells.

In this work, we studied the expression and sub-cellular localization of NMDARs in developing cultured hippocampal neurons by a combined approach of imaging and molecular techniques. The present results indicate that NMDAR subunits are expressed and localized in the soma, minor neurites, and axons during neural polarization. More importantly, at these early stages, NMDARs are functional since NMDAR agonists increased Ca^2+^ influx from the extracellular space, whereas specific antagonists blocked this response. The loss of function of NMDARs delayed neuronal polarization and decreased axonal elongation, whereas their gain of function promoted neuronal polarization and axonal development.

Additionally, previous studies indicate that hippocampal neurons release glutamate spontaneously into the extracellular milieu early in development (Andreae, Ben Fredj, and Burrone, 2012; Andreae and Burrone, 2015), before synaptic contact formation. Therefore, the role of glutamate as a diffusible factor that controls neuronal morphology early in development merits exploration. The present results support the hypothesis that glutamate acts as a diffusible factor, which, through NMDARs, promotes early neuronal development. Of note, NMDAR activation promoted Ca^2+^-induced Ca^2+^ release (CICR) from the Endoplasmic Reticulum (ER) through Ryanodine (RyR) and inositol 1,4,5-trisphosphate (IP_3_) receptors (IP_3_R) channels^2^). In addition, NMDARs modulated Rho GTPase Rac1 activity and actin cytoskeleton dynamics at the axonal growth cone and regulated the physiological levels of ROS. These findings suggest that there is a link between glutamatergic signaling and NOX2 complex activity that, through the dual function of Rac1, supports neuronal polarity and axonal development. These results provide a novel regulation layer connecting NMDAR functions in the axon with Ca^2+^ release from the ER and Rac1 activation of polarizing neurons.

## Materials and Methods

### Hippocampal primary cultures

Dissociated hippocampal neuron cultures were prepared from embryonic (18.5) rats as described previously (Kaech and Banker, 2006; Wilson et al., 2020) but without a feeding glial layer. The Bioethical Research Committee of the Universidad de Chile and the ANID manual for animal experimentation approved the protocols, so all animal experiments were carried out according to these protocols. Briefly, the hippocampi were dissected in Hank’s balanced salt solution (HBSS) supplemented with HEPES, incubated for 20 min at 37°C with trypsin (0,25% final concentration), and then washed with HBSS. The hippocampi were mechanically dissociated in 10% HS MEM. After that, neurons were counted and seeded on Poly-L-Lysine (1 mg/mL) coated glass coverslips (or in Poly-L-Lysine coated plastic dishes). Cell culture density was established according to each specific experiment. Once the neurons were attached (1 hour after plating), the 10% HS MEM was replaced with Neurobasal^TM^ Medium supplemented with B-27TM, GlutaMAX^TM^, sodium pyruvate, and Penicillin-Streptomycin antibiotics (all from GIBCO). The neuronal cultures were maintained for up to 3 days in vitro at 37°C and 5% CO2 in a humidified atmosphere. It is worth mentioning that the Neurobasal^TM^ media and the B-27^TM^ supplement do not contain glutamate, and the GlutaMAX^TM^ reagent (which is not glutamate) is more suitable than L-glutamine for cell cultures. Both the cDNAs encoding NMDARs subunits (GluN1-GFP, GluN2A-GFP, and GluN2B-GFP) and shRNA-GluN2A and shRNA-GluN2B were kindly provided by Brigitte Van Zundert, Universidad Andres Bello, Chile (Luo et al., 2002; Kim et al., 2005; Sepulveda et al., 2010).

### Transfection

Neurons were transiently transfected with Lipofectamine 2000 (Life Technologies) in Neurobasal medium according to the manufacturer’s instructions. After 2 hours of transfection, neurons were supplemented with B-27, GlutaMAX, sodium pyruvate, and antibiotics. Experiments were performed 24-72 h after cDNA transfection.

### Ca^2+^ imaging

Neurons (1.5 x 10^2) were cultured in 35 mm dishes on 25 mm glass coverslips pretreated with Poly-L-Lysine (1 mg/mL). After 1 DIV or 3 DIV, neurons were loaded with Fluo4-AM (5 µM) (Life Technologies) for 20 min in Hank’s balanced salt solution supplemented with HEPES (HBSS) at 37°C and 5% CO2. Then, the Fluo4-AM-containing medium was replaced with a previously recovered neurobasal medium for 40 minutes under the same incubation conditions. Later, the coverslips were mounted into a video microscopy chamber with HBSS or Krebs’ solution containing the following (in mM): 126 NaCl, 2.5 KCl, 25 NaHCO_3_, 1.2 NaH_2_PO_4_, 1.2 MgCl_2_, and 2.5 CaCl_2_. Time-lapse images were acquired every 2 s for 5 min with a confocal microscope using 20X magnification. To induce NMDAR-mediated Ca^2+^ influx, the neurons were stimulated with RSTG (10 µM) after 1 minute of baseline fluorescence. Regions of interest (ROIs) were defined at the soma of neurons. The average baseline fluorescence intensity was defined as "F0”, whereas “F” represents the fluorescence intensity after stimulation. The ratio F/F0 was used to determine the fold-change of Fluo4-AM fluorescence intensity after stimulation. The analyses were performed using the "time series analyzer" plug-in from Fiji-ImageJ (NIH, Bethesda).

### Antibodies

The following primary antibodies were used: GluN1 (mouse, Merck Millipore; 1: 500 for immunoblotting and 1: 200 for immunofluorescence), GluN2A, and GluN2B (rabbit, Thermo; 1:500 for immunoblotting and 1: 250 for immunofluorescence), α-tubulin (mouse, Sigma; 1:10.000 for immunoblotting), Tau-1 (mouse, Merck Millipore; 1: 5.000 for immunoblotting and 1:500 for immunofluorescence), β3-tubulin (mouse, Promega; 1: 1.000 for immunofluorescence), MAP2 (rabbit, Merck Millipore; 1:500 for immunofluorescence), GFP (rabbit, Covance 1:100 for immunofluorescence) and Alexa Fluor 405-, 488-, 543- or 647-conjugated secondary antibodies (1:400; Life Technologies), Alexa Fluor 647-conjugated Phalloidin (1:140, Invitrogen).

### Protein extracts and immunoblotting

Hippocampal neurons (1.5 x 10^6^ cells/well) were cultured in 35 mm dishes pretreated with poly-L-lysine (1 mg/ml) for 1 or 2 DIV (for stage 2 and 3 neurons, respectively). They were lysed in lysis buffer (150 mM NaCl, 50 mM Tris-HCl [pH 8.0], 1% Triton X-100, 0.5% sodium deoxycholate, 0.1% SDS, 1 mM PMSF, 1 mM NaF, 100 nM Calyculin A) and proteases inhibitors cocktail (Roche, Applied Science). The lysates were centrifuged at 14,000 rpm for 20 minutes, the supernatant was denatured, and then immunoblotting was performed. The membranes were incubated with primary antibodies diluted in 1% nonfat milk dissolved in 0.05% TBST overnight at 4°C with agitation. After washing the membranes with 0.05% TBST, they were incubated with suitable HRP-conjugated secondary antibodies for 90 minutes at room temperature. Proteins were detected using the Pierce ECL Western Blotting Substrate.

### Immunofluorescence cell staining

Neurons were fixed with 4% paraformaldehyde (PFA) in phosphate-buffered saline (PBS) containing 4% sucrose for 25 minutes at 37°C. After washing with PBS, cells were permeabilized with 0.2% Triton X-100/PBS for 5 minutes. The cells were then blocked with 5% bovine serum albumin (BSA)/PBS for 1 hour at room temperature and incubated with each indicated antibody overnight at 4°C. After washing, samples were incubated with the appropriate Alexa Fluor-conjugated secondary antibody for 45 minutes at room temperature, then mounted using Fluor Save™ Reagent (Merck Millipore).

### Extracellular glutamate detection

Neurons were cultured (320 neurons/mm²) in 24 multi-well plates on glass coverslips pretreated with poly-L-lysine (1 mg/mL). We maintained them for 1, 2, or 3 days in vitro (DIV) in Neurobasal Medium minus phenol red (to avoid autofluorescence), supplemented with B27, Glutamax, sodium pyruvate, and antibiotics. Notably, none of these solutions contained glutamate (Brewer et al., 1993; Chen et al., 2008) (Invitrogen). Conditioned media samples (50 µL out of 500 µL) were collected from neuronal cultures at 1, 2, or 3 DIV before neuron fixation. Samples were centrifuged at 10,000 rpm for 5 minutes at 4°C to remove possible cellular debris, and then kept at −80°C until glutamate detection. The ultrasensitive Glutamate Assay Kit (ab252893, Abcam) was used to detect and quantify the neurotransmitter glutamate from conditioned media according to the manufacturer’s instructions. This method is based on enzymatic reaction and fluorescence detection (Ex/Em=535/587) and produces a strong signal proportional to glutamate quantity. The kit detects specific glutamate at very low concentrations (minimum sensitivity of 0.05 µM). Enzymatic glutamate detection was performed in kinetic mode (37°C for 1 hour) using the Synergy 2 multiple-detection microplate reader (Biotek). Glutamate was detected and quantified using a standard curve with known concentrations (from 0 to 10 µM).

### FRET measurements

Neurons were cultured (4 x 10^4 cells/well) in 24 multiwell plates on glass coverslips pretreated with poly-L-lysine (1 mg/mL). After 1 DIV, neurons were transiently transfected with the Rac1 FRET-2G biosensor to measure Rac1 activity. Image acquisition and FRET efficiency analysis were performed as described previously (Wilson et al., 2015).

### Actin dynamics analysis

Neurons were cultured (1.5 x 10^5 cells/well) in 35 mm dishes on 25 mm glass coverslips pretreated with poly-L-lysine (1 mg/mL). After 1 DIV, neurons were transiently co-transfected with the Lifeact-GFP biosensor and either shRNA control, shRNA-GluN2A, or shRNA-GluN2B cDNA. After 24 hours of expression, the coverslips were mounted into a video microscopy chamber with the neurons’ growth medium. Time-lapse images were acquired every 15 seconds for 10 minutes with a confocal microscope using 40X magnification.

### H_2_O_2_ detection

Neurons were cultured (4 x 10^4 cells/well) in 24 multi-well plates on glass coverslips pretreated with poly-L-lysine (1 mg/mL). After 1 DIV, neurons were transiently transfected with the HyPer2 ratiometric biosensor to detect local intracellular H_2_O_2_ production (Markvicheva et al., 2011). The transfected neurons were excited at 405 nm and 488 nm and the emission at 505-530 nm was collected. For the HyPer H_2_O_2_ analysis, the H_2_O_2_ content was measured by dividing the fluorescence emission recorded following excitation at 488 nm by the fluorescence emission recorded at excitation 405 nm (Belousov et al., 2006; Wilson et al., 2015).

### Image acquisition and analysis

Immunofluorescence, Ca^2+^ imaging, FRET, and HyPer experiments were performed using an LSM 710 confocal microscope (Zeiss). All experiments were analyzed using Fiji-ImageJ (NIH, Bethesda).

### Statistics

Data are presented as the mean ± SEM of at least three independent cultures. For statistical significance between two groups, the Student’s t-test was used, and one-way ANOVAs to compare three or more groups. All analyses were performed using GraphPad Prism 8 software.

## Results

### NMDARs are expressed and functional during neuronal polarity acquisition of cultured hippocampal neurons

Growing evidence has described the expression of the glutamate NMDA receptor subunits at different developmental stages (Washbourne, Bennett, and McAllister, 2002; Washbourne et al., 2004; Song et al., 2009; Wang et al., 2011; Bustos et al., 2014). However, NMDAR subunit expression during the process of neuronal polarization and axonal growth remains elusive. Therefore, the endogenous expression and cellular localization of NMDARs subunits in immature cultured hippocampal neurons were evaluated (Fig. 1). By immunoblotting, the expression of GluN1, GluN2A, and GluN2B subunits of NMDARs in stage 2 (non-polarized) and 3 (polarized) cultured neurons (1 DIV and 2 DIV, respectively) (Fig. 1A) was detected. Furthermore, western blot assays showed that all subunits here studied increased their expression in cultured hippocampal neurons from 1 DIV to 2 DIV (Fig. 1A). Next, the subcellular distribution of these subunits using an immunofluorescence approach was evaluated in stage 2 and stage 3 cultured hippocampal neurons. As illustrated in Fig. 1B-D, GluN1, GluN2A, and GluN2B subunits were present in non-polarized (stage 2) and polarized (stage 3) neurons. They displayed a punctate and clustered distribution throughout the neurons and were distributed in the soma and minor neurites in stage 2 neurons (Fig. 1B-D); interestingly, they were distributed along the axonal compartment, including the growth cone structure in stage 3 neurons (Fig. 1B-D). Together, these data show that NMDARs are expressed early in development, including during neuronal polarization stages.

**Figure 1.**
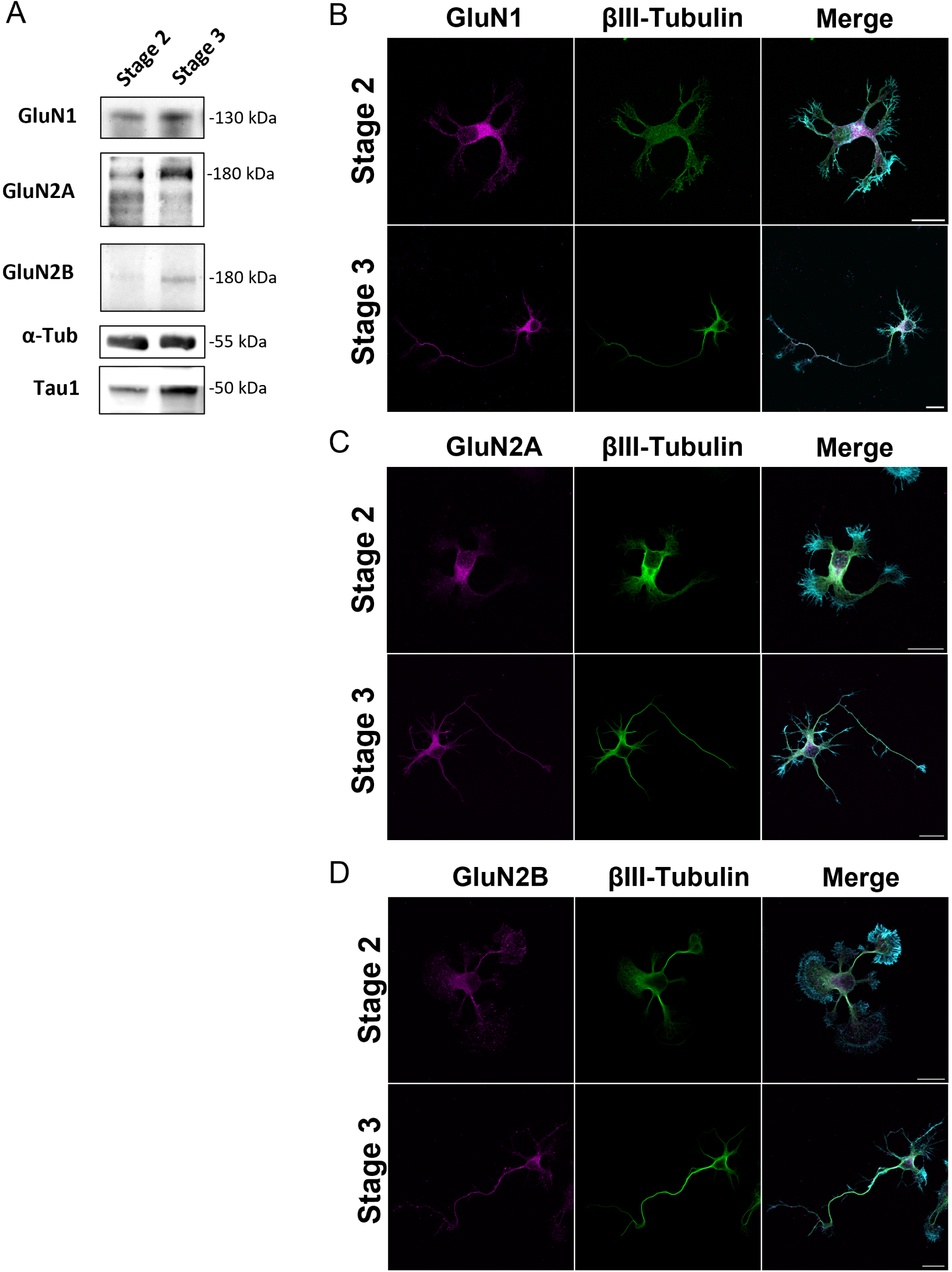
NMDARs expression and sub-cellular localization during the acquisition of neuronal polarity. **A**, Detection by immunoblotting of GluN1, GluN2A, and GluN2B subunits at stage 2 and stage 3 cultured neurons, 24 h and 48 h, respectively. **B - D**, Sub-cellular localization of GluN1, GluN2A, and GluN2B subunits in neurons at stage 2 and stage 3, 24 h and 48 h, respectively. NMDAR subunits were detected both at soma and minor neurites in stage 2 cultured neurons and across the axon of stage 3 cultured neurons. Scale bar: 20 μm.

Next, the expression and subcellular localization after overexpression of GFP-tagged NMDARs subunits (Luo et al., 2002) during neuronal polarization was evaluated (Fig. 2). To this aim, neurons were co-transfected immediately after plating with vectors coding RFP and GFP-tagged NMDARs subunits and fixed at 1 and 2 DIV. To enhance NMDAR detection in neurons, an anti-GFP antibody was used according to previous protocols in the field (Luo et al., 2002; Prybylowski et al., 2002; Washbourne, Bennett and McAllister, 2002; Bresler et al., 2004; Wang et al., 2011; Gill et al., 2014; Willems et al., 2020). Of note, GFP-tagged GluN1, GluN2A, and GluN2B subunits were distributed in the soma of stage 2 neurites and in the axons of polarized neurons (stage 3) and were especially evident in axonal growth cones (Fig. 2A-C). These results align with the endogenous expression of NMDAR subunits shown in Figure 1, reinforcing that ectopic expression of NMDARs resembles native immunodetection of NMDARs during polarity acquisition.

**Figure 2.**
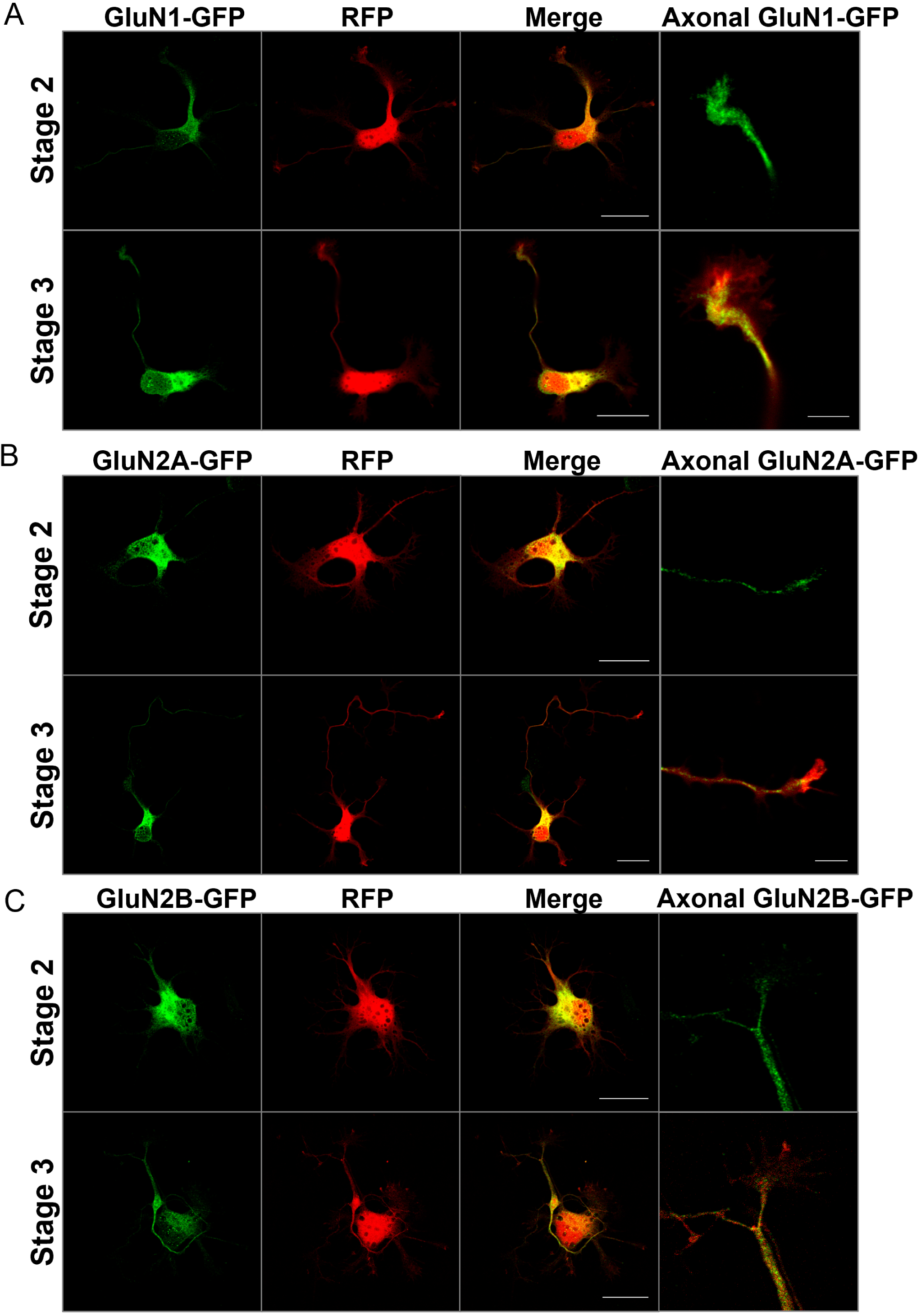
Sub-cellular localization of transfected GFP tagged-NMDARs subunits during neuronal polarization. **A-C**, Neurons were cultured and co-transfected immediately after plating with GluN1-GFP, GluN2A-GFP, or GluN2B-GFP and RFP constructs and fixed at 1 or 2 DIV to evaluate the NMDARs subunits distributions in stage 2 and 3 neurons. Anti-GFP antibody was used to amplify the GFP signal of GFP-tagged-NMDARs subunits. Scale bar: 20 μm.

Since functional NMDARs form heterotetrameric complexes that enable Ca^2+^ influx into neurons (Paoletti, Bellone, and Zhou, 2013), NMDAR functionality by live-cell Ca^2+^ imagingwas then assessed. To this aim, neurons were loaded with Fluo4-AM (a Ca^2+^-sensitive fluorescent probe) to measure cytoplasmic Ca^2+^ levels in neurons challenged with the specific NMDARs agonist (RS)-Tetrazol-5-yl) glycine (RSTG) (Schoepp et al., 1991; Furukawa et al., 2005). Imaging was performed through time-lapse recording and RSTG stimulation was done at t=1 min until completing recording for 5 min (Figure 3A). Treatment with RSTG increased cytoplasmic Fluo4 fluorescence; this increase did not occur in Ca^2+^-free HBSS medium (Fig. 3A and B), indicating that RSTG induces Ca^2+^ influx into the neuron from the extracellular medium. Next, neurons were treated with either D-AP5 (100 µM), a competitive NMDAR antagonist that prevents glutamate binding to the NR2A and NR2B subunits (Evans et al., 1982), or with MK-801 (5 µM), a non-competitive channel blocker (Wong et al., 1986; Huettner and Bean, 1988). Both D-AP5 and MK-801 inhibited Ca^2+^ influx induced by RSTG (Fig. 3A and C), suggesting that an NMDAR-mediated mechanism contributes to the development of yet unpolarized neurons (stage 2). Similarly, treatment of polarized neurons (stage 3) with RSTG increased Fluo4-AM fluorescence (Fig. 3D and E); this increase did not take place in Ca^2+^-free medium (Fig. 3D and E) and after treatment with D-AP5 or MK-801 (Fig. 3D and F), suggesting that NMDARs are active in early polarized neurons. Moreover, the amplitude of the Ca^2+^ response was higher in stage 3 than in stage 2 neurons, suggesting a developmental-dependent phenotype, which is consistent with the protein levels of NMDARs shown in Fig. 1A. Together, these data suggest that immature polarizing neurons express functional NMDARs that modulate Ca^2+^ influx into neuronal cells.

**Figure 3.**
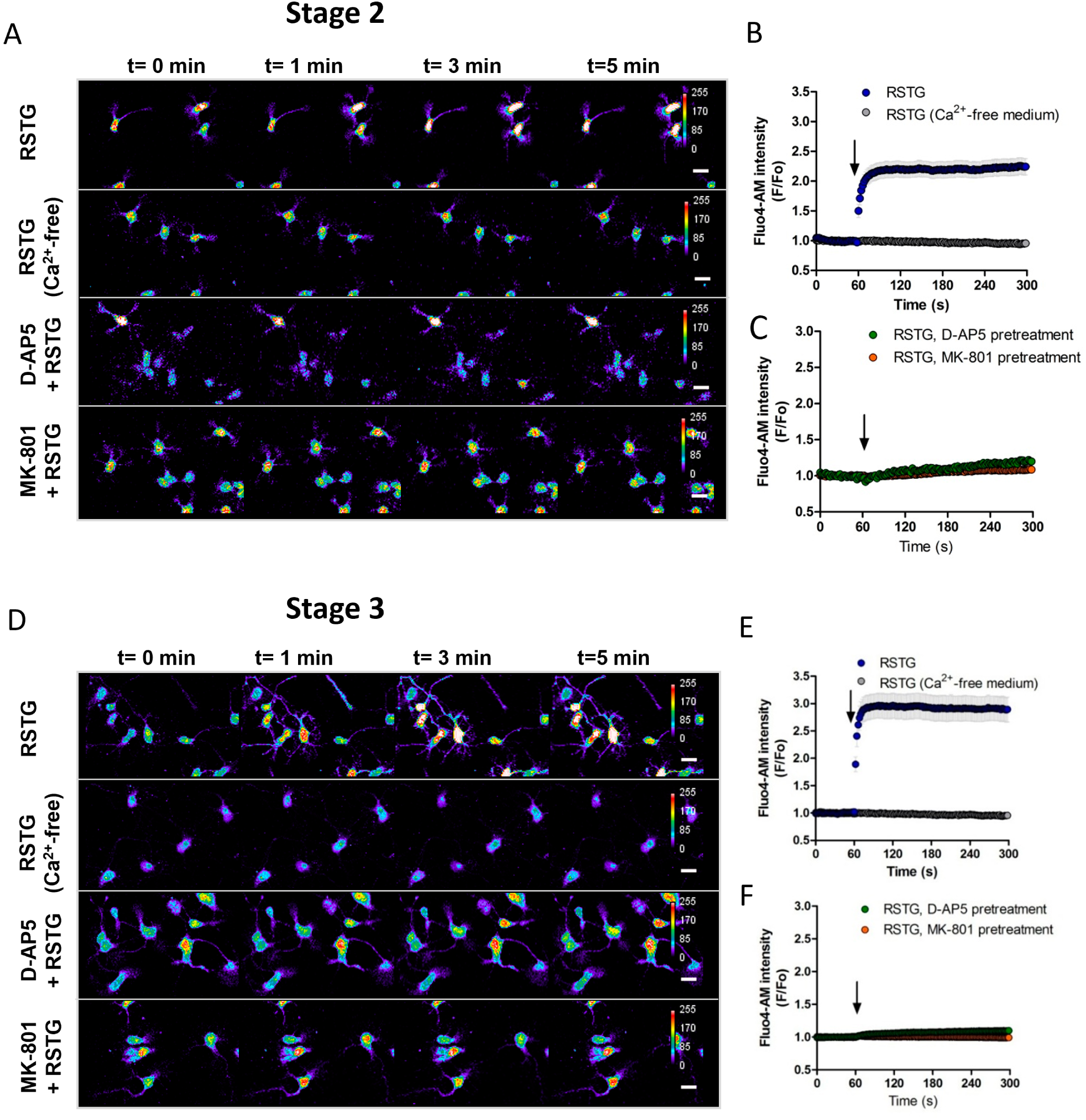
The NMDARs are functional during the acquisition of neuronal polarity. Neurons at 1 and 3 DIV were loaded with Fluo-AM to visualize cytoplasmic Ca2+ in live neurons. **A** and **D**, Representative time-lapse imaging of the responses to the agonist RSTG (10 μM), to RSTG in Ca^2+^-free medium and in neurons pre-treated with the NMDARs inhibitors D-AP5 (100 μM) or MK-801 (5 μM) for 1 h before agonist addition in stage 2 and 3 neurons, respectively. **B**, Fluo4-AM fluorescence after RSTG stimulation in Ca^2+^-containing or Ca^2+^-free medium during the recording of Ca^2+^ signals illustrated in **A**. **C**, Fluo4-AM fluorescence after RSTG stimulation in neurons pre-treated with the inhibitors D-AP5 or MK-801 of **A**. Scale bar: 20 μm.

### The NMDARs contribute to the establishment of hippocampal neuronal polarity and axonal growth

The NMDARs are commonly associated with glutamatergic synaptic transmission (Hunt and Castillo, 2012). However, they are also instrumental for the regulation of dendritic branching and axonal elongation in mature neurons, uncovering functions beyond synapses (Schmitz et al., 2009; Sepulveda et al., 2010; Bustos et al., 2014). However, the role of NMDARs during polarity acquisition has yet to be properly characterized. In this regard, our data showed that, at this timeframe, neurons express functionally active NMDARs. Thus, we tested whether NMDARs contribute to regulate neuronal polarity acquisition. To this aim, we used pharmacological and genetic approaches to evaluate the contribution of NMDARs to axonal specification and elongation. Firstly, neurons were treated with D-AP5 or MK-801 (NMDARs inhibitors) or NMDA or RSTG (NMDAR agonists) at 1 DIV, when neurons remained unpolarized and were fixed at 3 DIV to evaluate axonal growth (Fig. 4 A-D). NMDAR inhibition induced a significant increase in the number of stage 1 and 2 neurons, to the detriment of the stage 3 population (Fig. 4A and B). Moreover, neurons treated with NMDAR inhibitors displayed reduced axonal growth, whereas NMDAR activation promoted axonal extension (Fig. 4A, C, and D). These results suggest that the activity of NMDAR promotes the polarization and axonal growth of developing neurons. Moreover, the AMPA receptor inhibitor CNQX (10 µM) did not affect polarity acquisition or axonal elongation (Fig. 4B and C), suggesting that glutamatergic signaling mediated by NMDARs shapes neuronal morphology.

**Figure 4.**
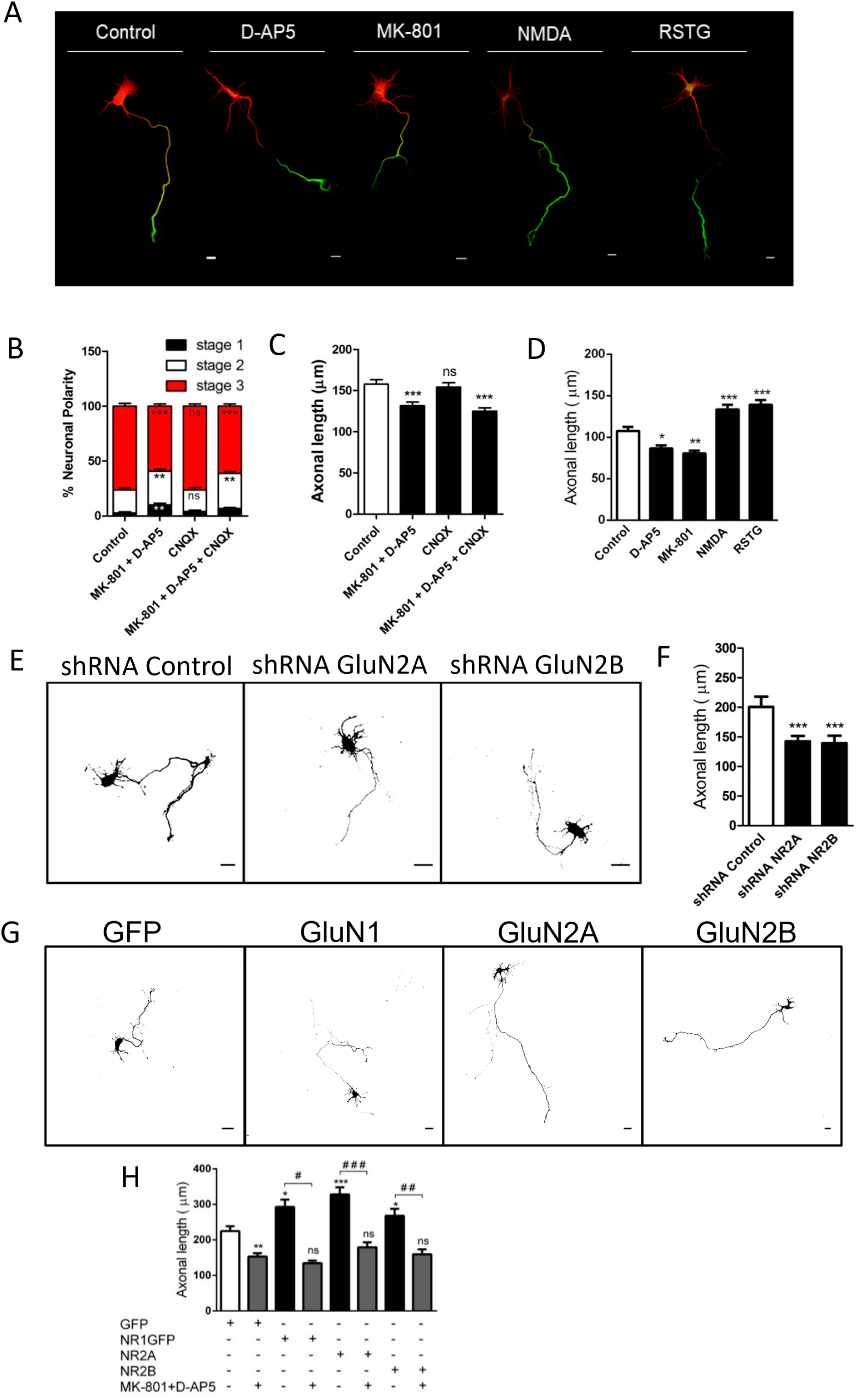
The NMDARs contribute to the establishment of hippocampal neuronal polarity and axonal growth. **A-D**, Neurons at 1 DIV were cultured and treated with the NMDARs inhibitors D-AP5 (100 μM) and/or MK-801 (5 μM) and with the agonist NMDA (50 μM) or RS-Tetrazol-Glycine (RSTG) (10 μM) and fixed at 3 DIV. Moreover, neurons were treated at 1 DIV with the AMPA inhibitor CNQX (10 μM) and fixed at 3 DIV. **A**, 3 DIV neurons were stained with MAP2 and Tau in the control condition after NMDARs inhibition or activation. **B-D**, Quantification of neuronal polarization and axonal growth of hippocampal neurons after the indicated treatments. **E-F**, Neurons were cultured and transfected immediately after plating with GFP with shRNA control or transfected with GFP with the shRNA-2A or shRNA-2B constructs and fixed at 3 DIV. **E**, Representative transfected neurons at 3 DIV. **F**, Quantification of axonal growth of panel E. **G** and **H**, Neurons were cultured and transfected immediately after plating with GFP alone (control) or co-transfected with the NMDARs subunits construct and fixed at 3 DIV. On the other hand, 1 DIV transfected neurons were treated with the inhibitor D-AP5 (100 μM) and MK-801 (5 μM) and were fixed at 3 DIV. **G**, Representative transfected neurons at 3 DIV. **H**, Quantification of axonal of panel G. Scale bar: 20 μm.

The contribution of NMDARs to neuronal polarization was examined next, using genetic manipulation. Neurons were transfected with plasmids encoding either shRNAs or wild-type cDNAs, for loss and gain of function assays, respectively. In this regard, we used plasmids previously validated in the literature to knock down GluN2A or GluN2B subunits (Kim et al., 2005). To visualize neurons expressing these constructs, a GFP-encoding plasmid was co-transfected as a reporter, and the results were compared with those displayed by neurons expressing a control shRNA sequence (shRNA-Luciferase). Neurons transfected after plating and fixed at 3 DIV for morphological analysis displayed restrained axonal growth following GluN2A or GluN2B silencing (Fig. 4E-F). These results align with the pharmacological blockade shown in this work, reinforcing that NMDAR inhibition halts polarization and axon extension in developing neurons.

Conversely, overexpression of GluN1, GluN2A, and GluN2B subunits significantly promoted axonal growth, which is especially evident in neurons expressing the GluN2A subunit (Fig. 4G-H). Furthermore, treatment with the inhibitors MK-801 or D-AP5 reversed this effect, supporting the specificity of NMDARs into the axon growth displayed by polarizing neurons (Fig. 4H). These data highlight the fact that functional NMDARs are required to promote neuronal polarity acquisition in cultured neurons and suggest that glutamate might be present in the extracellular milieu, supporting early neuronal development. Thus, these results reopen the debate on the extra-synaptic functions of NMDARs, particularly in their role in the polarization and axon specification of neurons.

### Extracellular detection of the neurotransmitter glutamate promotes axonal development

Previous reports suggest that polarized hippocampal neurons (4-5 DIV) release glutamate spontaneously into the extracellular space, contributing to dendritic remodeling (Andreae et al., 2012; Andreae and Burrone, 2015). However, the presence of extracellular glutamate during the acquisition of neuronal polarity has not been described. We then sought to evaluate the presence of glutamate in the extracellular space and its endogenous effect on neuronal development through two complementary approaches.

First, by using an ultrasensitive and specific Glutamate Assay Kit, we directly evaluated the presence of glutamate in the extracellular milieu, by collecting conditioned medium from 1-3 DIV hippocampal neuronal cultures (Fig. 5). We detected glutamate concentrations of 0.85, 0.81 and 1.21 µM in 1, 2 and 3 DIV cultures, respectively. Measurements were blanked using a neurobasal maintenance medium in the absence of neurons, suggesting the release of glutamate during neuronal polarity acquisition (Fig. 5A). At 3 DIV, when neurons are fully polarized, extracellular glutamate levels were the highest within the timeframe analyzed; these findings are consistent with other reports describing the spontaneous release of glutamate from the axonal compartment at this stage (Andreae et al., 2012). Interestingly, at 1 DIV, when neurons remain in the unpolarized stage 2, we found a significant glutamate increase, suggesting its release not only before synapse formation but also before the break in neuronal symmetry, consistent with our data on the expression and functionality of NMDARs in non-polarized neurons.

**Figure 5.**
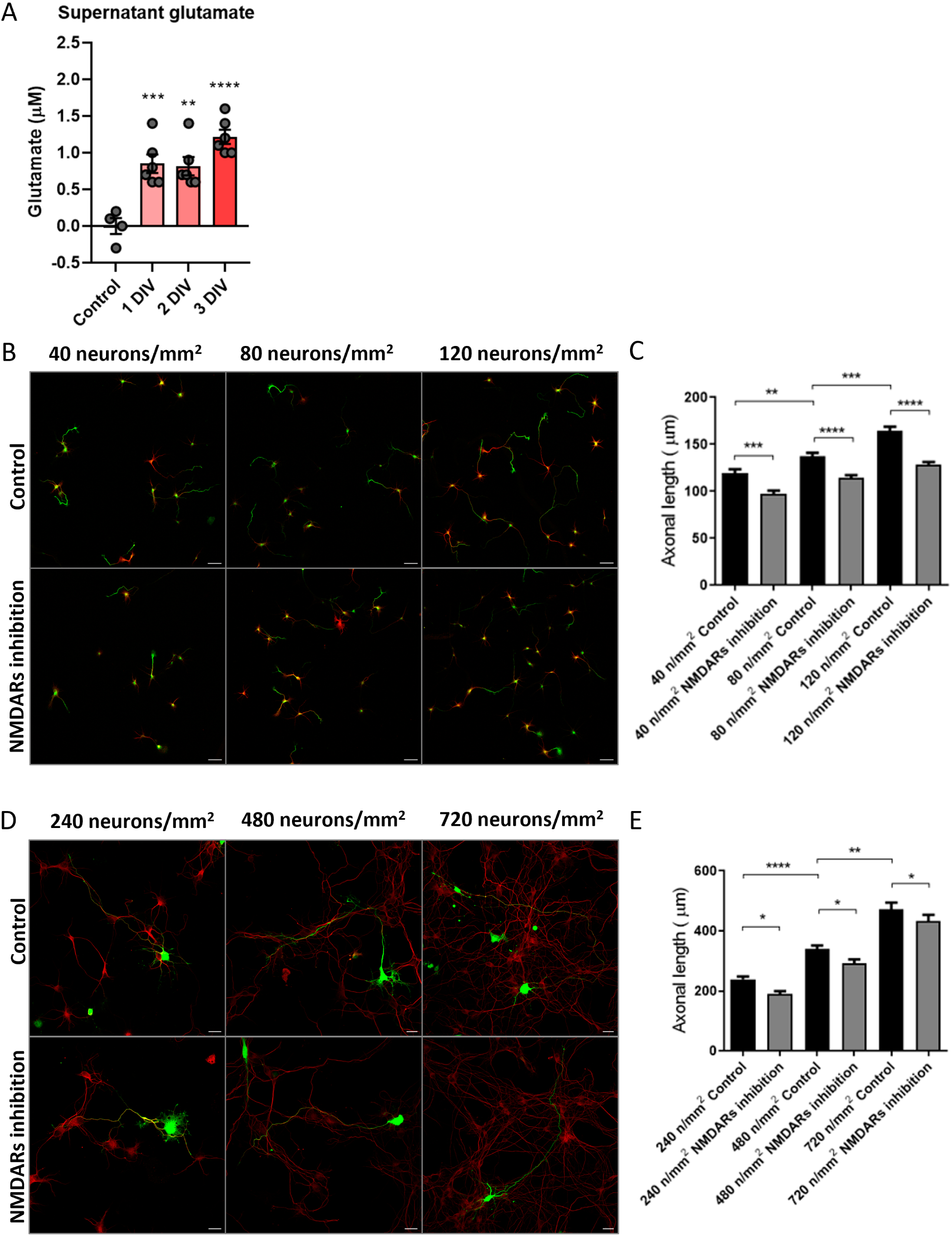
Physiological glutamate promotes NMDAR activation to support axonal elongation. **A**, Supernatants from 1, 2, or 3 DIV neuronal cultures (320 neurons/mm2) were recovered to detect the glutamate neurotransmitter throughout neuronal development specifically. We detected the presence of extracellular glutamate from 1, 2, and 3 DIV, while it was not detected in the medium without neurons (control), suggesting glutamate release from neurons. **B-C**, 1 DIV neurons cultured at increasing densities (40 to 120 neurons/mm^2^) were treated with vehicle or the NMDAR inhibitors D-AP5 (100 μM) and MK-801 (5 μM) and then fixed at 3 DIV. **A**, Representative 3 DIV neurons stained with MAP2 and Tau in control and after treatment of NMDARs inhibitors. **C**, Quantification of axonal growth of hippocampal neurons from panel B. Scale bar: 50 µm. **D-E**, Neurons cultured at increasing densities (240 to 720 neurons/mm2) were transfected immediately after plating with GFP and treated at 1 DIV with NMDARs inhibitors D-AP5 (100 μM) and MK-801 (5 μM) and then fixed at 3 DIV. **D**, Representative GFP transfected 3 DIV neurons and stained with βIII-tubulin (red) in control and NMDARs inhibition conditions. **E**, Quantification of axonal growth of hippocampal neurons from panel D. Scale bar: 20 µm.

Second, since cell density in neuronal cultures modifies axon elongation rate and neuronal development, the presence of glutamate and the contribution of NMDARs to axonal growth was evaluated at different neuronal densities (ranging from 40 to 720 neurons per mm²; Fig. 5B-E). We hypothesized that the higher the neuronal confluency *in vitro*, the higher the release of glutamate stimulates axonal growth. Of note, neither the Neurobasal medium nor its supplements contain glutamate (Brewer et al., 1993; Chen et al., 2008). To this end, we seeded neurons under a progressive confluency culture scheme, ranging from low densities (40, 80, and 120 neurons per mm²) to high densities (240, 480, and 720 neurons per mm^2^). At 1 DIV, neurons were treated with control vehicle (H_2_O) or NMDARs inhibitors (D-AP5 or MK-801) and fixed at 3 DIV to evaluate their morphology and axonal elongation by MAP2 and Tau1 immunofluorescence (Fig. 5B). Within the low range, axonal extension paralleled the increase in cell density, which was abolished in the presence of NMDAR inhibitors (Fig. 5B-C). Next, higher cell densities were used (240, 480, and 720 neurons per mm2), and axonal elongation was evaluated. To visualize single neurons, as needed for precise morphological analyses in high-density cultures, neurons were transfected after plating with a GFP-encoding plasmid and treated with vehicle or NMDAR inhibitors after 1 DIV. Then, neurons were fixed at 3 DIV to evaluate neuronal morphology and axonal growth. The results of these treatments reproduced the trend observed in the low-density range (Fig. 5D-E). Together, these results indicate that neurons release glutamate spontaneously into the extracellular space by acquiring neuronal polarity and axonal outgrowth, promoting axonal growth via NMDARs activation.

### NMDAR activation promotes Ca^2+^ release from the Endoplasmic Reticulum during neural polarization

Having shown the participation of glutamate and NMDARs in neural polarization and axonal growth, we explored the intracellular molecular mechanisms activated by NMDARs that regulate neuronal morphology. First, we focused on the role of Ca^2+^ ions, which permeate through activated NMDARs, functioning as a second messenger. In addition, Ca^2+^ ions have been previously associated with axon elongation (Henley and Poo, 2004; Zheng and Poo, 2007; Wilson et al., 2016). Second, Ca^2+^ influx from the extracellular medium promotes Ca^2+^ release from internal Ca^2+^ stores by a mechanism termed Ca^2+^-induced Ca^2+^ release (CICR) (Fabiato and Fabiato, 1975; Hidalgo and Arias-Cavieres, 2016). In mature neurons, NMDAR-mediated Ca^2+^ influx triggers CICR by activating inositol 1,4,5-trisphosphate receptor (IP_3_R) and Ryanodine receptor (RyR) channels (Verkhratsky and Shmigol, 1996; Berridge, 1998), which are expressed in the Endoplasmic Reticulum (ER) and are essential for neuronal polarity and axonal elongation (Wilson et al., 2016). However, NMDAR-dependent CICR during neuronal polarization remains understudied. To test its role in neuronal polarization, intracellular recordings of Ca^2+^ levels in non-polarized and polarized primary neurons were performed (Fig. 6). Stage 2 or Stage 3 neurons were loaded with Fluo4-AM and incubated for 1 h before Ca^2+^ recordings with the IP_3_R inhibitor 2-APB (10 µM) (Ruiz et al, 2009) or ryanodine at an inhibitory concentration (25 µM) to block RyR channels (Adasme et al., 2015; Wilson et al., 2016; More et al., 2018). Figure 6A shows representative Ca^2+^ signals in Stage 2 neurons before (0-1 min) and after (1-5 min) NMDAR stimulation. Treatment with 2-APB or inhibitory ryanodine reduced the fluorescence intensity elicited by RSTG (Fig. 6A-C), suggesting that Ca^2+^ entry mediated by NMDARs promotes Ca^2+^ release through IP_3_R and RyR channels in non-polarized neurons. Similarly, in polarized neurons, the fluorescence intensity induced by NMDAR stimulation was significantly reduced following IP_3_R or RyR inhibition (Fig. 6D-F). Together, these data suggest that Ca^2+^ influx through NMDARs promotes Ca^2+^ release from the ER, mediated by IP_3_R or RyR channels during neuronal polarity acquisition. Also, these results confirm that NMDARs present in non-polarized and polarized neurons are functional and coupled to the regulation of intracellular Ca^2+^ levels.

**Figure 6.**
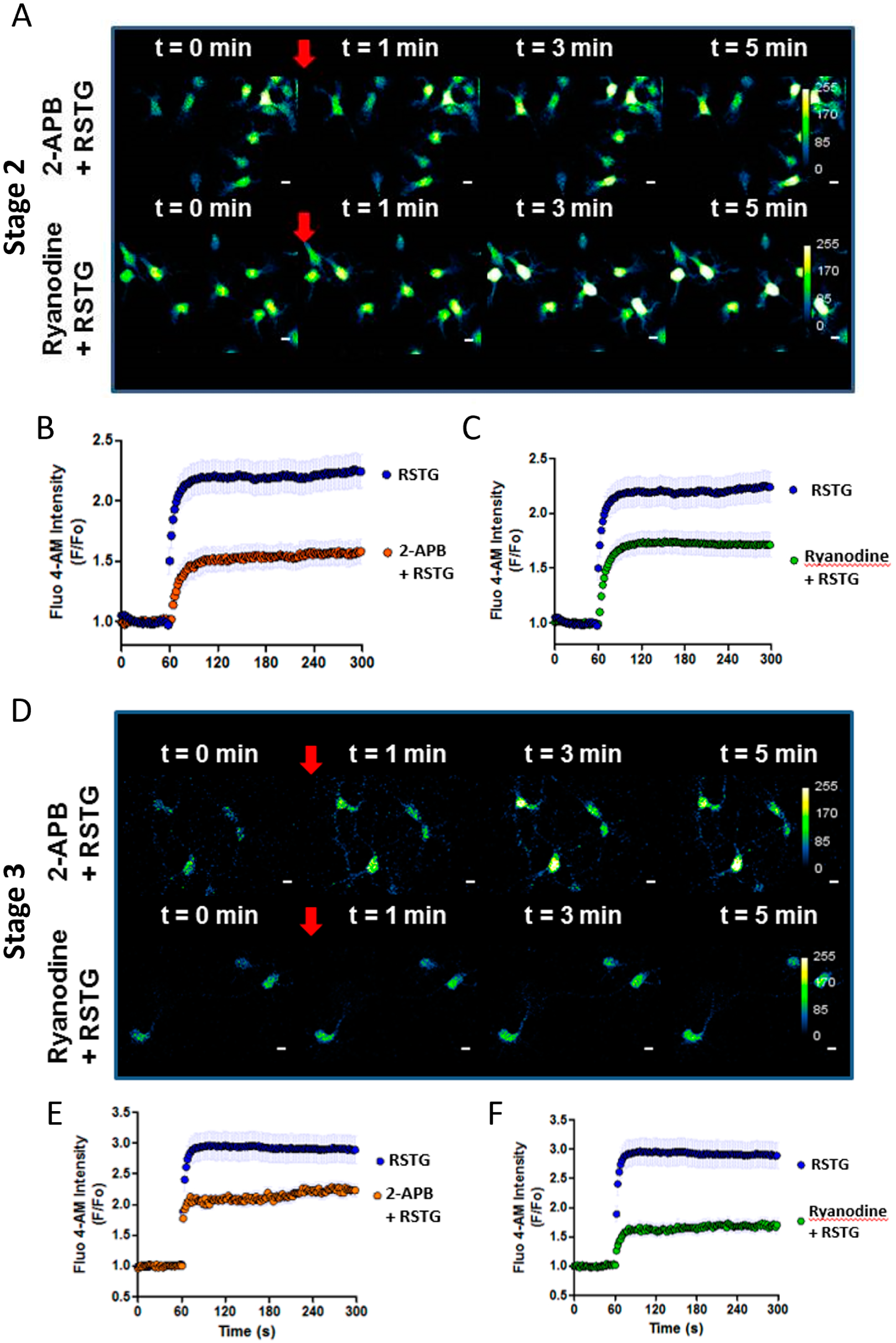
NMDAR activation promotes Ca^2+^ release from the Endoplasmic Reticulum. **A** and **D**, 1 and 3 DIV neurons were loaded with Fluo4-AM to visualize cytoplasmic Ca^2+^ in live neurons. **A** and **D**, Representative time-lapse imaging for stage 2 and 3 neurons pre-treated with IP3R inhibitor 2-APB (10 μM) or with ryanodine (25 μM) for 1 h to inhibit ryanodine receptors before agonist addition, respectively. **B-C** and **E-F** Fluo4-AM fluorescence after RSTG stimulation in stage 2 and 3 neurons pre-treated with the inhibitors 2-APB or ryanodine of panels A and D, respectively. Scale bar: 20 μm.

### NMDARs regulate Rac1 activity and actin cytoskeleton remodeling during neuronal polarization

Previous reports indicate that the increase in cytoplasmic free Ca^2+^ concentration derived from extracellular Ca^2+^ influx and intracellular Ca^2+^ release regulates the activity of the small monomeric protein Rho GTPase Rac1 (Fleming et al., 1999; Price et al., 2003; Jin et al., 2005). The Rac1 protein is a master regulator of the actin cytoskeleton and controls different aspects of neuronal development (Govek, Newey and Van Aelst, 2005; Tahirovic et al., 2010; Gonzalez-Billault et al., 2012; Wilson et al., 2016). Moreover, NMDARs activation induces Rac1 activation in dendrites of olfactory system neurons (Aihara et al., 2021). However, the contribution of NMDARs in regulating Rac1 and actin cytoskeleton dynamics during neuronal polarization has not been reported. To address this issue, stage 2 neurons were transfected with the FRET-2G probe to evaluate the activity of Rac1 (Fig. 7) (Martin et al., 2016). After 1 day of NMDAR inhibition, the polarized neurons were fixed to evaluate FRET efficiency using confocal microscopy. We found that neurons treated with NMDAR inhibitors showed a decrease of Rac1 FRET efficiency, either in the whole neuron or the axonal compartment (axon), suggesting that NMDAR activity regulates Rac1 activity (Fig. 7A-C). Of note, NMDAR inhibition restrained again axonal growth and neuronal polarization, reinforcing its contribution in maintaining neuronal outgrowth (Fig 7A).

**Figure 7.**
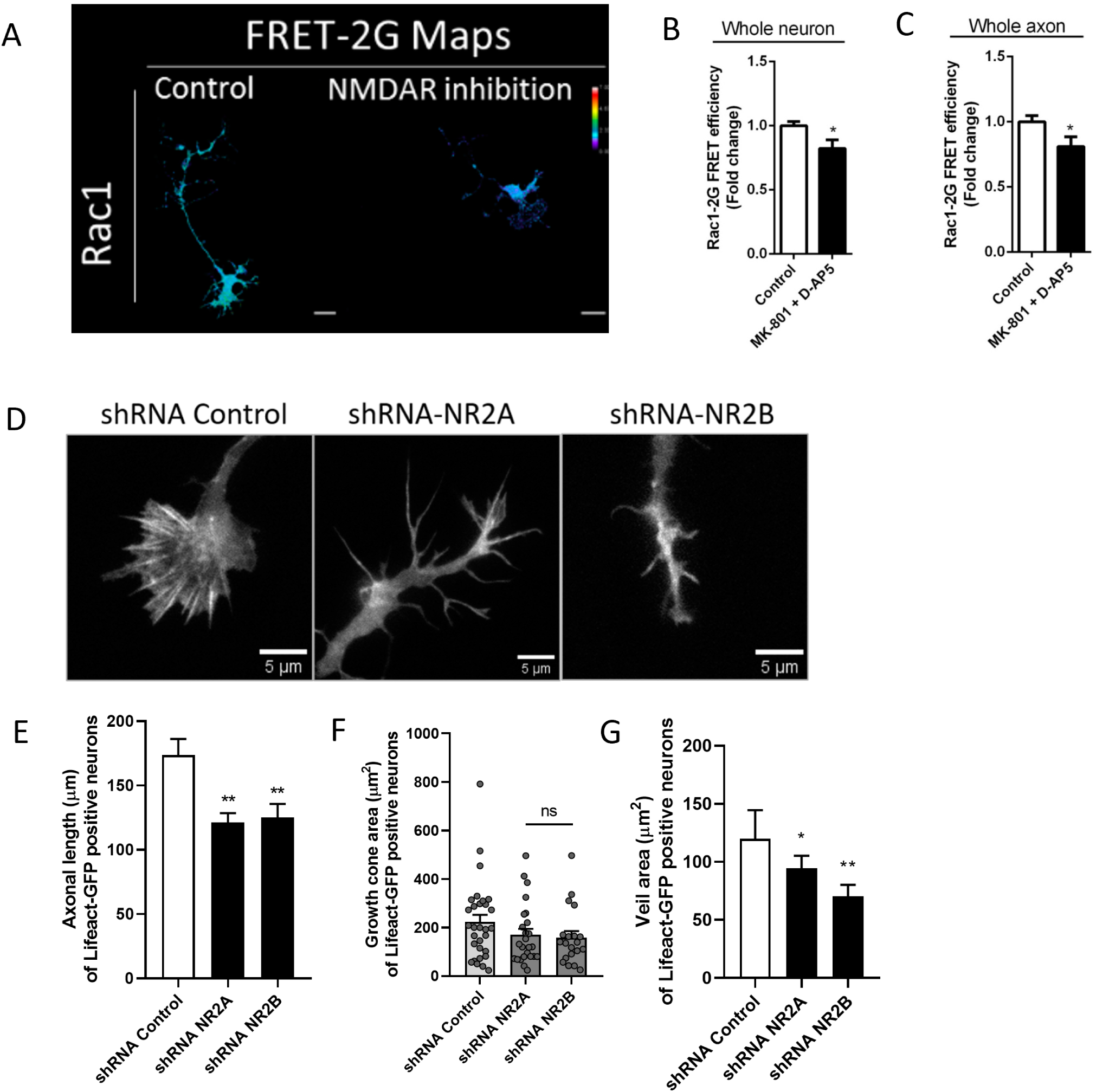
NMDARs pharmacological and genetics inhibition regulate actin dynamics and Rac1 activity. **A-C**, Hippocampal neurons of 1 DIV were transfected with the Rac1-2G probe to measure the FRET efficiency after NMDAR inhibition (D-AP5 and MK-801) during FRET-2G probe expression. **A**, FRET-2G maps to Rac1. **B** and **C**, Quantification of the Rac1 FRET efficiency in control and NMDARs inhibitors-treated neurons in whole neuron and axonal compartment, respectively. **D**, Representative axonal growth cone of neurons expressing Lifeact-GFP in shRNA control, shRNA-GluN2A, or shRNA-GluN2B neurons to visualize F-actin structure. **E**, Quantification of the axonal length of Lifeact-GFP positive neurons. F, Quantification of the area of axonal growth cone. G, Quantification of axonal growth cones’ lamellar structures (veil area). Scale bar: 5 μm.

Rac1 regulates the formation of axonal lamellipodia structure and its dynamics (Tahirovic et al., 2010; Sayyad, Fabris, and Torre, 2016). Therefore, actin organization using the genetically encoded probe Lifeact-GFP was evaluated next (Riedl et al., 2008). Because NMDAR inhibition decreased axonal elongation, we focused our analysis on the axonal growth cone (axonal tip) to evaluate its morphology as a proxy for actin polymerization and axonal outgrowth. To this aim, stage 2 neurons were co-transfected with Lifeact-GFP along with shRNA control or shRNAs targeting GluN2A and GluN2B subunits (Fig. 7D-G). As expected, silencing each regulatory subunit significantly reduced axonal growth compared with the control (Fig. 7D-E) as previously observed (Fig. 4E-F), confirming that both regulatory subunits contribute to axonal growth.

We observed a fan-shape structure at the tip of axons, corresponding to growth cone with characteristic shape, including lamellar structures and filopodia. In contrast, neurons treated with shRNAs targeting NR2A or NR2B exhibited collapsed growth cones (Fig. 7D). Additionally, we observed a wide variation in the area of the growth cones (Fig. 7F), consistent with previous reports in the literature (Bagonis et al., 2019). We evaluated the total area (lamellipodia + filopodia) of growth cones of neurons in control and under knock-down conditions; no differences were found after silencing the NMDARs (Fig. 7F). However, suppressing NMDARs significantly reduced lamellar structures (veil area), which aligns with the decay of Rac1 activity after NMDAR inhibition (Fig. 7G). These data indicate that basal NMDAR activity is needed to sustain Rac1 activity and the dynamics of the actin cytoskeleton which supports axonal growth during neuronal polarization.

### NMDARs contribute to H_2_O_2_ production in early development

Physiological ROS levels are needed to support and promote different aspects of neuronal development and functionality (Munnamalai and Suter, 2009; Olguín-Albuerne and Morán, 2015; Wilson et al., 2015; Beckhauser et al., 2016; Bórquez et al., 2016; Wilson et al., 2016; Wilson et al., 2018; Muñoz-Palma and González-Billault, 2021). In addition, previous reports suggest a link between glutamatergic signaling and superoxide production in cortical mature neurons (Brennan et al., 2009; Reyes et al., 2012; Brennan-Minnella et al., 2013). However, coupling between NMDAR activity and ROS production during neuronal polarization remains unreported. To examine this issue, stage 2 neurons were transfected with the HyPer2 biosensor (Markvicheva et al., 2011) to evaluate intracellular H_2_O_2_ levels using confocal microscopy in polarizing neurons (Wilson et al., 2015; Wilson et al., 2016) (Fig. 8). At stage 3, neurons were treated with the NMDAR agonist RSTG and fixed 5 or 15 min after stimulation, which induced an increase in H_2_O_2_ intracellular levels analyzed in the whole neuron and the axonal compartment (Fig. 8A-C). Next, H_2_O_2_ levels were evaluated in the presence of NMDAR inhibitors (Fig. 8D-F). Neurons expressing HyPer2 were treated with D-AP5 and MK-801 during 24 h and then fixed. NMDAR inhibition decreased the H_2_O_2_ levels measured in the whole neurons and the axonal compartment (Fig. 8D-F). Interestingly, NMDAR inhibition for 1 h significantly reduced H_2_O_2_ levels (Fig. 8D-F), suggesting that NMDARs actively contribute to H_2_O_2_ production. We then examined ROS production after NMDARs loss of function. To this aim, stage 2 neurons were co-transfected with the HyPer2 biosensor along with either shRNAs control or targeting the GluN2A or GluN2B subunits, and fixed at 2 DIV for imaging (Fig. 8G-I). GluN2A and GluN2B silencing significantly decreased H_2_O_2_ levels measured in the whole neuron and the axonal compartment (Fig. 8H-I), confirming our previous pharmacological experiments. Therefore, these data suggest that NMDAR activity contributes to regulating H_2_O_2_ production during neuronal polarization.

**Figure 8.**
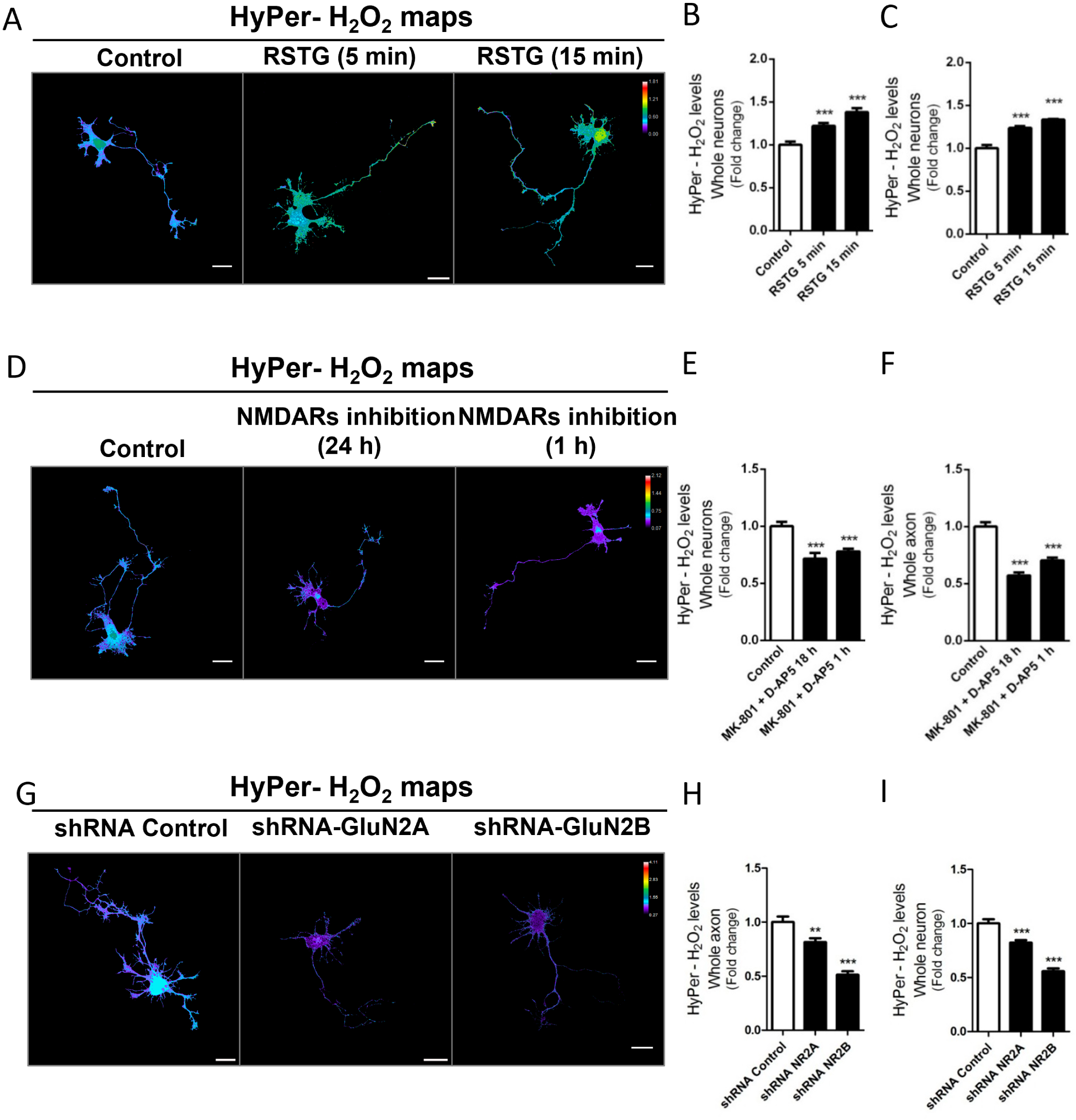
NMDARs regulate ROS production during the acquisition of neuronal polarity. **A-I**, 1 DIV Neurons were transfected with the HyPer2 biosensor to measure H_2_O_2_ local output. **A-C**, After 1 day of expression, neurons were treated with the agonist RSTG (10 μM) for 5 and 15 min. **A**, Representative H_2_O_2_ map of both control and NMDAR activation. Quantification of the HyPer- H_2_O_2_ levels in whole neuron (**B**) and axon (**C**) from neurons of panel A. D, Representative HyPer- H_2_O_2_ maps of both control and neurons treated with NMDARs inhibitors (D-AP5 and MK-801) for one day and one h. Quantification of the HyPer- H2O2 levels in whole neuron (**E**) and axon (**F**) from neurons of panel D. G-I, 1 DIV Neurons were transfected with the HyPer2 biosensor together with the shRNA control, shRNA-2A or shRNA-2B constructs and fixed at 2 DIV. **G**, Representative H_2_O_2_ maps of the different conditions. Quantification of the HyPer- H_2_O_2_ levels in whole neuron (H) and axon (I) from neurons of panel G. Scale bar: 20 μm.

### Dual function of Rac1 during neuronal polarization: regulation of actin cytoskeleton and NOX2-mediated H_2_O_2_ production

Since inhibition of NMDARs reduced the activity of Rac1, a protein involved in both actin remodeling and H_2_O_2_ production via the NOX2 complex, we then focused on evaluating this dual function of Rac1 during neuronal polarization. To probe this hypothesis, we used 2 strategies to split Rac1 functions. On one hand, the inhibitor Phox-I1 was used to block the specific interaction between Rac1 and p67phox (Bosco et al., 2012), which is essential for NOX2 activation. Consequently, we evaluated the H_2_O_2_ content in stage 2 neurons expressing the HyPer2 sensor, treated for 24 h with the Phox-I1 inhibitor (Fig. 9A-B). Blocking the interaction between Rac1 and p67phox significantly reduced the intracellular H_2_O_2_ content (Fig. 9A-B).

**Figure 9.**
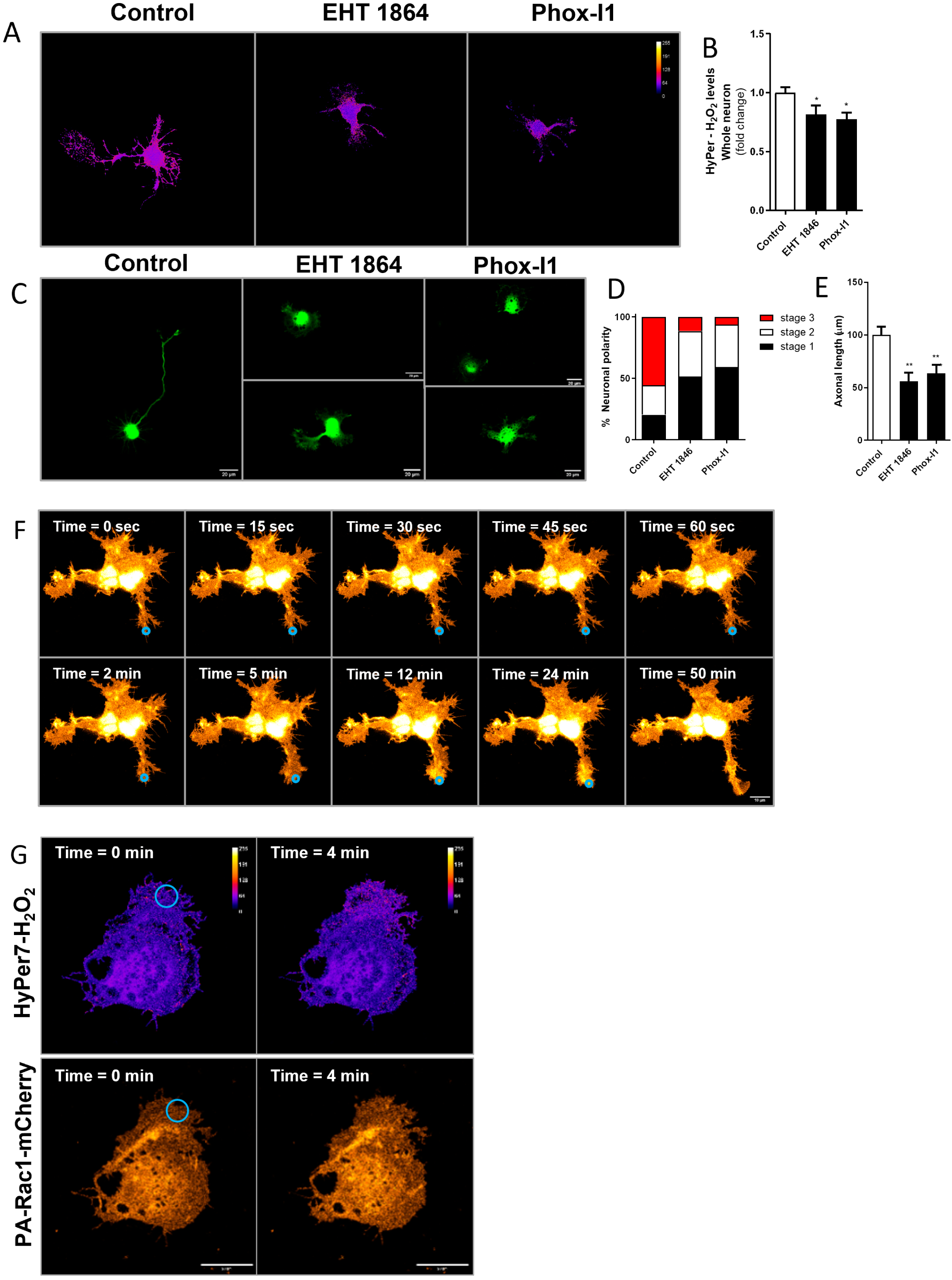
Rac1 regulates NOX2-mediated H_2_O_2_ production during the acquisition of neuronal polarity. **A**, 1 DIV neurons were transfected with the HyPer2 biosensor to measure local H_2_O_2_ and were treated with EHT-1964 (total Rac1 inhibitor) or Phox- I1 (Rac1-p67phox interaction inhibitor). **B**, Both EHT-1864 and Phox-I1 reduced the intracellular H_2_O_2_ content. Scale bar: 20 μm. **C**, Neurons were transfected immediately after plating with GFP, treated at 1 DIV with EHT-1864 or Phox-I1 inhibitors, and then fixed at 3 DIV. **D-E**, Both EHT-1864 and Phox-I1 reduced neuronal polarization and axonal length. F, Hippocampal neuron was transfected with the Photoactivatable Rac1-Q61L-mCherry (PA-Rac1). Persistent local illumination with 458 nm light at the growth cone of minor neurite rapidly generates lamellipodia protrusion, which results in neurite extension. Scale bar: 10 μm. **G**, Local illumination at the axonal growth cone of developing neuron co-expressing PA-Rac1 and HyPer7 rapidly generates both lamellipodia protrusion and H_2_O_2_ production, demonstrating that Rac1 activation leads to actin cytoskeleton dynamics and H_2_O_2_ production. Scale bar: 20 μm.

Next, we measured H_2_O_2_ content in neurons treated with EHT-1864, a specific inhibitor of the Rac1 isoform, which binds to the GDP/GTP binding site and thus interrupts the on-off cycling of the GTPase (Shutes et al., 2007; Onesto et al., 2008). Of note, EHT-1864 reduced the H_2_O_2_ content to a similar extent as PhoxI-1 (Fig. 9A-B), suggesting that basal Rac1 activity regulates H_2_O_2_ production through NOX2. Then, neuronal morphology was evaluated after treatment with PhoxI-1 and EHT-1864 inhibitors. Neurons were transfected with GFP after plating, treated with these inhibitors at 1 DIV, and then fixed at 3 DIV for morphological analysis. We found that both Rac1 inhibitors delayed neuronal polarization and reduced axonal length (Fig. 9C-E), suggesting that Rac1 activity and its assembly into the NOX complex are necessary to promote neuronal development.

Finally, we sought to evaluate the simultaneous cellular effects promoted by optogenetic Rac1 activation. To this aim, neurons were transfected with the Photoactivatable Rac1-Q61L-mCherry (PA-Rac1) version (Wu et al., 2009). Stage 2 neuron expressing PA-Rac1 was activated by 458 nm illumination at the growth cone of a minor neurite. Local and persistent activation generated lamellipodia protrusion, which resulted in neurite extension, showing that Rac1 activation promotes neurite outgrowth locally (Fig. 9F). We then sought to evaluate H_2_O_2_ production after PA-Rac1 activation. Local illumination at the axonal growth cone rapidly generated both lamellipodia protrusion and H_2_O_2_ production (Fig. 9G), demonstrating the dual function of Rac1 as a promoter of actin cytoskeleton dynamics and H_2_O_2_ production during neuronal polarization.

## Discussion

### NMDARs expression during neuronal polarity acquisition

Our results provide new insights and strong evidence supporting the unexpected idea that NMDARs play essential roles during early neuronal development and axonal outgrowth. Traditionally, NMDARs have been exclusively associated with neurotransmission and their location is assumed to be the postsynaptic terminal (Daw, Stein, and Fox, 1993). Nevertheless, emerging evidence suggests that NMDARs exhibit a broader distribution, beyond postsynaptic locations. Several reports indicate their presence within the axonal compartment and presynaptic terminals in several neurons, suggesting pivotal roles of NMDARs in supporting neurotransmission from the presynaptic side (Zheng, Wan and Poo, 1996; Corlew et al., 2008, 2008; Schmitz et al., 2009; Rodriguez-Moreno et al., 2011; Wang et al., 2011; Buchanan et al., 2012; Gill et al., 2014; Wong et al., 2021). In this study, our results unveil the expression, localization, and functionality of NMDARs across developing hippocampal neurons *in vitro*. Significantly, our findings underscore that both native and overexpressed NMDAR subunits are early expressed in development and are distributed within the axon and the axonal growth cone, particularly during the crucial steps of neuronal polarity acquisition. Consistent with our findings, similar results have been reported in already polarized hippocampal and cortical neurons (Washbourne, Bennett, and McAllister, 2002; Washbourne et al., 2004; Song et al., 2009; Wang et al., 2011, 2018).

However, certain discrepancies have surfaced concerning the axonal localization and exclusion of NMDARs in developing neurons. These disparities also extend to their involvement of NMDARs in presynaptic terminals during neuronal transmission (Wong et al., 2021). Several reports have highlighted divergencies on axonal distribution of native and ectopic expression of GFP-NMDARs in hippocampal neurons. For instance, the presence of GFP-GluN2B subunits in the axonal compartment at 3 DIV, disappearing by 5 DIV was reported (Song et al., 2009). By contrast, the axonal localization of GluN2B at 6 DIV, followed by a decay at 20 DIV was also reported (Herkert, Röttger, and Becker, 1998).

Furthermore, other studies have reported the presence of native GluN1 subunits and GFP-tagged GluN2A and GluN2B subunits in the axonal growth cone at 6 days *in vitro* (DIV) followed by a significant decrease in the localization of transfected YFP-GluN1, GFP-GluN2A, and GFP-GluN2B subunits at the distal axon in 16 DIV neurons (Wang et al., 2011), whereas the native GluN1 subunit was detected in axonal growth cone at 10 DIV (Ehlers et al., 1998). These discrepancies may arise from several factors, including difficulties in generating specific antibodies against NMDAR subunits, fluorescence detection methods, and challenges in detecting endogenous and ectopic expression of NMDARs in neurons, which often require further signal amplification, presenting an additional layer of complexity. Therefore, current evidence strongly indicates that NMDA receptors are expressed early and localized in the axons of developing neurons; once neurons reach functional maturation, they are restricted to the dendrites. Moreover, other studies suggest that NMDARs expression and localization are developmentally regulated (Corlew et al., 2008; Rodriguez-Moreno et al., 2011; Wang et al., 2011; Buchanan et al., 2012; Gill et al., 2014; Wong et al., 2021), moving past the controversy. Thus, the presence of NMDAR subunits in the axonal compartment and the axonal growth cone early in development suggests that NMDARs could play essential roles in early neuronal polarization and axonal outgrowth events.

### Role of NMDARs during the early developmental stage

In addition to their well-established roles in neuronal transmission, learning, and memory, extensive literature underscores the significance of NMDARs in various facets of neuronal development. Thus, NMDARs not only are involved in dendritic branching, spine formation, and maturation (Bustos et al., 2014; Mu et al., 2015) but they are also involved in neuronal migration (Komuro and Rakic, 1993; Behar et al., 1999; Kumada and Komuro, 2004) and neuritogenesis (Pearce, et al., 1987; Rashid and Cambray-Deakin, 1992). The results presented here indicate that NMDARs contribute to neuronal polarity acquisition and axonal growth of young hippocampal neurons. These findings agree with previous reports showing that NMDARs mediate axonal growth cone turning in cultured Xenopus spinal neurons (Zheng, Wan, and Poo, 1996), the promotion of axonal elongation, and growth cone splitting of dopaminergic neurons (Schmitz et al., 2009) in already polarized neurons. Moreover, Ca^2+^ as a second messenger, regulates neurite outgrowth and axonal growth cone motility (Henley and Poo, 2004; Zheng and Poo, 2007). Therefore, it is plausible that Ca^2+^ influx through NMDARs contributes to promote axonal growth, especially considering the axonal distribution of NMDARs described in this and other works. Whereas inhibition of NMDARs reduced dendritic dynamics and growth (Rajan and Cline, 1998; Rajan et al., 1999), GluN2A and GluN2B overexpression promotes dendritic arborization (Ewald et al., 2008; Sepulveda et al., 2010). Our results align with this view, since pharmacological and genetic inhibition of NMDARs restrained axonal elongation, suggesting that the NMDAR subunits GluN2A and GluN2B are functionally expressed and support axonal growth.

In contrast, overexpression of NMDAR subunits promoted axonal extension. Additionally, the contribution of GFP-GluN2A to axonal growth was more significant than those of GluN1 and GluN2B. The differential expression of these subunits and NMDAR stoichiometry may underlie these results (Bellone and Nicoll, 2007; Tárnok et al., 2008; Paoletti et al., 2013; Bustos et al., 2014).

### Functional NMDARs during the stage of neuronal polarization

The heterotetrameric nature of NMDARs requires their traffic towards the plasma membrane to form the channel pore, allowing Ca^2+^ influx when activated, and thus supporting NMDARs functionality (Barria and Malinow, 2002; Luo et al., 2002; Traynelis et al., 2010; Paoletti et al., 2013). Here, we showed that NMDARs are functional in the presence of their agonists during the acquisition of neuronal polarity. Under physiological conditions, the activation of NMDARs requires simultaneously the presynaptic release of glutamate and postsynaptic membrane depolarization (Mayer et al., 1984; Nowak et al., 1984; Espinosa and Kavalali, 2009). Interestingly, our Ca^2+^ imaging experiments revealed that, in addition to being inhibited by D-AP5, NMDARs were also inhibited by MK-801, a dose-dependent open-channel blocker of NMDARs (Wong et al., 1986; Huettner and Bean, 1988; Atasoy et al., 2008; McKay et al., 2013; Song et al., 2018). Accordingly, early in development, NMDARs may be activated through mechanisms in different ways for their traditional role as postsynaptic coincidence detectors, which typically require specific forms of activation (Wang et al., 2011; Wong et al., 2021). However, this possibility has yet to be thoroughly investigated.

### Glutamate and functional NMDARs during early stages of neuronal polarity acquisition

Finally, the evidence obtained from intracellular Ca^2+^ recordings indicates that NMDARs are functionally active early in development and shape neuronal morphology. Interestingly, the fact that the MK-801 blocks the NMDAR channel pore in a use-dependent manner (Wong et al., 1986; Huettner and Bean, 1988; Atasoy et al., 2008; McKay et al., 2013; Nabavi et al., 2013), supports our findings, suggesting physiological activation of NMDARs. Therefore, the physiological presence of glutamate could activate NMDARs early in development, prior to synaptic establishment. Glutamate release from the presynaptic terminal has been described to occur both in an evoked and spontaneous manner in mature neurons (Ramirez and Kavalali, 2011; Kavalali, 2015). However, previous reports suggest that spontaneous glutamate release occurs in young hippocampal neurons (Andreae et al., 2012; Andreae and Burrone, 2015). Of note, we detected the presence of glutamate in the extracellular milieu during neuronal polarization and observed a cell density dependence on axonal growth sensitive to NMDAR inhibitors.

### Intracellular signaling mediated by NMDARs during neuronal polarization

Because the contribution of NMDARs to neuronal polarity and axonal growth was unknown until now, the molecular mechanisms involved remained cryptic. In this regard, the CICR mechanism has been described to occur in mature neurons, mostly sustained by voltage-gated Ca^2+^ channels and NMDARs (Verkhratsky and Shmigol, 1996; Berridge, 1998). Our work supports this view and shows that CICR also occurs in polarizing neurons, mostly maintained by NMDAR-dependent signaling. Moreover, Ca^2+^ influx mediated by NMDARs and ER Ca^2+^ release regulates the activity of Rho GTPases in several cell types (Li, Van Aelst, and Cline, 2000; Sin et al., 2002; Govek et al., 2005; Jin et al., 2005). Our results show that NMDARs regulate Rac1 activity, a molecular determinant for axonal growth (Gonzalez-Billault et al., 2012). Moreover, their activity also impacts on the actin cytoskeleton of the axonal growth cone, most-likely supporting axonal outgrowth early in development. Additionally, the notion that H_2_O_2_ production mediated by the NOX2 complex regulates neuronal polarity, and neurite and axonal outgrowth has recently emerged (Olguín-Albuerne and Morán, 2015; Wilson, Núñez and González-Billault, 2015; Wilson et al., 2016; Wilson, Muñoz-Palma and González-Billault, 2018; Muñoz-Palma and González-Billault, 2021). Consistent with H_2_O_2_ production mediated by NMDARs in mature neurons (Brennan et al., 2009; Reyes et al., 2012), our data showed that NMDARs regulate H_2_O_2_ production during early axonal development, suggesting a link between glutamatergic signaling and H_2_O_2_ production during neuronal polarity acquisition. As previously reported, our data indicate coupling between Ca^2+^ influx through NMDARs function and H_2_O_2_ production mediated by the NOX2 complex (Brennan et al., 2009). However, our data also suggest a new combination of molecular intermediates to maintain the functional coupling mentioned above, by considering the involvement of NMDARs-mediated Ca^2+^ influx, CICR and Rac1 activity in the regulation of both the actin cytoskeleton dynamics and NOX2-mediated H_2_O_2_ production (Munnamalai and Suter, 2009; Munnamalai et al., 2014; Wilson, Núñez and González-Billault, 2015; Wilson et al., 2016; Acevedo and González-Billault, 2018), supporting acquisition of neuronal polarity and axonal development.

In conclusion, the present findings show that NMDARs expressed early in development promote axonal elongation through a mechanism involving Ca^2+^ signaling, Rac1 activity, actin cytoskeleton dynamics, and H_2_O_2_ production mediated by the NOX2 complex, supporting the notion that glutamatergic signaling mediated by NMDARs are instrumental not only for neuronal maturation and neurotransmission but also for shaping neurons early in development.

## Acknowledgments

This work was supported by the following grants: ANID/Fondecyt #1220414 and ANID/FONDAP #15150012 to CG-B. Spelling, grammar, and syntax were improved using the Grammarly platform.

